# Engineered bacteria recruit and orchestrate anti-tumor immunity

**DOI:** 10.1101/2022.06.16.496462

**Authors:** Thomas M. Savage, Rosa L. Vincent, Sarah S. Rae, Lei Haley Huang, Alexander Ahn, Kelly Pu, Fangda Li, Courtney Coker, Tal Danino, Nicholas Arpaia

**Affiliations:** Department of Microbiology & Immunology, Columbia University, New York, NY; Department of Pathology and Cell Biology, Columbia University, New York, NY; Department of Biomedical Engineering, Columbia University, New York, NY; Herbert Irving Comprehensive Cancer Center, Columbia University, New York, NY; Data Science Institute, Columbia University, New York, NY

## Abstract

Tumors employ multiple mechanisms to actively exclude or suppress adaptive immune cells involved in anti-tumor immunity. Strategies focused on overcoming these immunosuppressive or exclusion signals – through localized delivery of chemokines that directly recruit immune cells into the tumor microenvironment – remain limited due to an inability to target therapeutics specifically to the tumor. Synthetic biology enables engineering of cells and microbes for tumor localized delivery, offering therapeutic candidates previously unavailable using conventional systemic administration techniques. Here, we engineer bacteria to produce and intratumorally release chemokines to attract adaptive immune cells into the tumor environment. Intravenous or intratumoral delivery of bacteria expressing an activating mutant of the human chemokine CXCL16 (hCXCL16^K42A^) leads to the recruitment of activated T cells within tumors and offers therapeutic benefit in multiple mouse tumor models. Furthermore, we rationally target an additional step in the immune activation cascade – specifically, the presentation of tumor-derived antigens by dendritic cells – using a second engineered bacterial strain expressing CCL20. This combined targeting approach led to the recruitment of type 1 conventional dendritic cells and effectively synergized with hCXCL16^K42A^-induced T cell recruitment to provide additional therapeutic benefit. In summary, we engineer bacteria to cooperatively recruit and activate both innate and adaptive anti-tumor immune responses, offering a new cancer immunotherapy strategy.

## INTRODUCTION

Surmounting the obstacles tumors employ to suppress immune cell infiltration has proven to be an elusive target. Immune infiltration, particularly by CD8^+^ cytotoxic T lymphocytes and, more specifically, memory CD8^+^ T cells, within tumors is widely considered a positive prognostic factor^1–4^. Immune cells undergo chemotaxis in response to chemokines^5,6^; however, although chemokines may be an attractive therapeutic target in cancer^7^, conventional drug delivery methods fail to generate tumor-localized chemokine gradients that sufficiently overcome immune cell exclusion^8,9^. Thus, despite the importance of tumor infiltration by immune cells in treatment outcomes, new approaches are needed to actively attract leukocytes into the tumor microenvironment.

Bacteria are increasingly recognized as a component of the tumor microenvironment, with genetic manipulations of bacteria enabling renewed consideration for bacterial cancer therapy^10–12^. A previously generated synthetic gene circuit in bacteria, termed the synchronized lysis circuit (SLC), enables repeated delivery of tumor-localized therapeutics following synchronized lysis^13–15^. Here, we use the SLC and engineer bacteria that produce tumor-localized chemokines to recruit innate and adaptive immune cells involved in both the priming and response phases of tumor immunity and activate a therapeutically effective anti-tumor immune response, a heretofore unavailable therapeutic approach.

## RESULTS

### Generation and characterization of CXCL16-variant strains in probiotic *E. coli*

The chemokine CXCL16 specifically recruits memory T cells with extra-lymphoid homing potential^16,17^, and its expression is associated with improved T cell infiltration and survival in colon and lung cancers, among other tumor types^18,19^. We therefore hypothesized that SLC-mediated release of CXCL16 in tumors would promote infiltration of activated T cells and support antitumor immunity (**Fig. 1A**). We first expressed the bioactive mature region (Asn30–Pro118) of human CXCL16 (hCXCL16) on a previously described^15^ high copy plasmid in the probiotic *E. coli* Nissle 1917 (EcN) and confirmed that release of hCXCL16 was SLC-dependent (**Fig. 1B**). Release of hCXCL16 in tumors *in vivo* also was SLC-dependent, with minimal detection in tumor homogenates from untreated tumors or those treated with EcN expressing hCXCL16 without SLC (-SLC), but significantly greater detection when SLC was present (+SLC) (**Fig. 1C**). To identify the optimal variant of CXCL16, we generated mouse CXCL16 (mCXCL16) and previously described^20^ activating (hCXCL16^K42A^) and inactivating (hCXCL16^R73A^) mutants of human CXCL16 (**Fig. 1D**). For functional assessment of the probiotic-derived CXCL16 variants, we developed a chemotaxis assay in which activated T cells were assayed for their migration in response to lysate of wildtype or variant-expressing EcN strains (**Fig. 1E**). Compared to wildtype lysate, activated mouse CD4^+^ and CD8^+^ T cells significantly migrated in response to lysates of EcN strains expressing mCXCL16 and activating hCXCL16^K42A^, but not wildtype hCXCL16 or inactivating hCXCL16^R73A^ (**Fig. 1F**). Consistent with previous characterization of the hCXCL16^K42A^ mutation^20^, activated human T cells displayed similar trends in response to lysate of the hCXCL16^K42A^-expressing EcN strain (**Fig. S1**). These data suggest that *E. coli*-derived hCXCL16^K42A^ potently attracts mouse and human activated T cells, potentially easing translational and pre-clinical studies, and warranting assessment of this strain *in vivo* using murine tumor models.

**Fig. 1.**
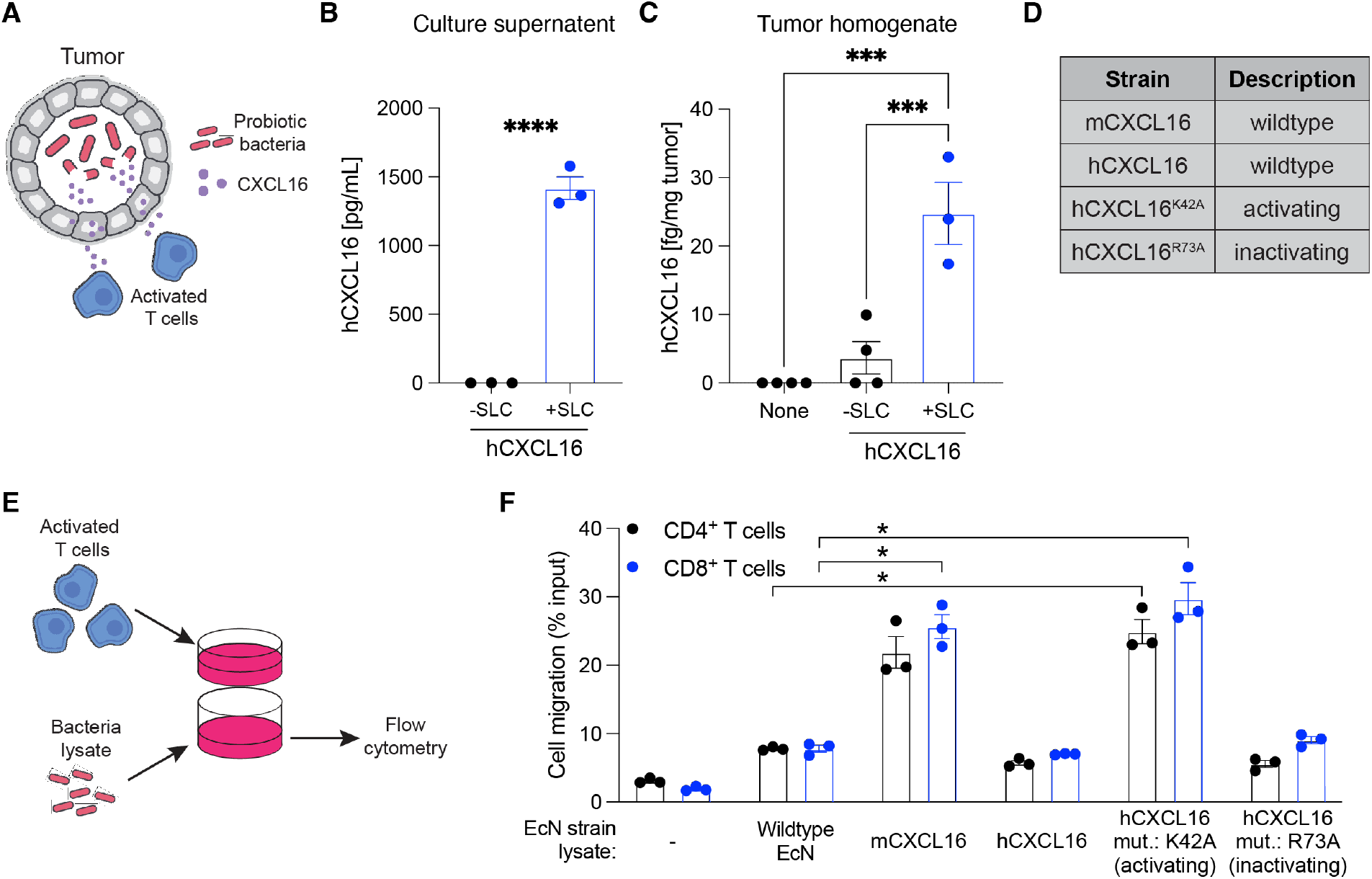
Generation and characterization of CXCL16-variant strains in probiotic *E. coli*. **(A)** Schematic overview of approach to recruit activated T cells to tumors via localized expression and release of CXCL16 by probiotic bacteria. **(B)** ELISA quantification of hCXCL16 concentration in culture supernatants of EcN expressing hCXCL16 with (“+SLC”), or without (“-SLC”), SLC. Data are representative of 3 independent experiments (\*\*\*\**P*<0.0001, 2-sided unpaired student’s t-test). **(C)** Quantification of hCXCL16 concentration in homogenates of A20 tumors intratumorally injected with EcN expressing hCXCL16 with (“+SLC”), or without (“-SLC”), SLC by ELISA. Data are representative of 2 independent experiments (*\*\*\*P*< 0.001, 1-way ANOVA with Holm-Sidak post-hoc test). **(D)** Table of CXCL16 variants expressed in EcN and used throughout the current study. **(E)** Schematic overview of chemotaxis assay used to assess the bioactivity of EcN- expressed CXCL16 variants. **(F)** Flow cytometric quantification of the migration of activated mouse T cells in response to each of the indicated EcN strain lysates. Media without any bacteria was used as a control Data are representative of 3 independent experiments (\**P*<0.05, 2-way ANOVA with Dunnett post-hoc test). All data displayed as mean ± s.e.m.

### Probiotic *E. coli*-derived CXCL16 promotes tumor regression

We examined the efficacy of EcN-derived hCXCL16^K42A^ *in vivo* by treating subcutaneously growing murine tumors after they were established and palpable (~100 mm^3^). We first tested variants of hCXCL16 in the A20 B cell lymphoma model, performing intratumoral injections of bacteria every 3–4 days for a total of 4 treatments (**Fig. 2A**). Significant reductions in tumor growth were observed in mice treated with SLC EcN strains expressing activating hCXCL16^K42A^ (eSLC-hCXCL16^K42A^) as compared to wildtype hCXCL16 (eSLC-hCXCL16)- and inactivating mutant hCXCL16^R73A^ (eSLC-hCXCL16^R73A^)-expressing SLC EcN strains (**Fig. 2B, S2A**), confirming the therapeutic activity of EcN-mediated intratumoral delivery of hCXCL16^K42A^ *in vivo*. Furthermore, the hCXCL16^K42A^ strain was significantly more effective in treating established A20 tumors than PBS or EcN expressing SLC alone (eSLC) (**Fig. 2C, S2B**). Most notably, the hCXCL16^K42A^ strain induced complete regression of 7 of 10 treated A20 tumors. Phenotyping of tumor infiltrating lymphocytes revealed that the hCXCL16^K42A^ strain induced an increase in activated and proliferating CD4^+^ conventional T (Tconv) cells, as assessed by Ki-67 expression and cytokine production (**Fig. 2D–2E, S2C**). Treatment with the hCXCL16^K42A^ strain also led to increased CD8^+^ T cell activation in A20 tumors, as assessed by Ki-67 (**Fig. 2D**), Granzyme-B expression (**Fig. 2F**), and their production of the cytokines IFNγ and TNFα following *ex vivo* restimulation with PMA and ionomycin (**Fig. 2G, S2D**). Moreover, upon *ex vivo* restimulation with an MHC-I (H-2K^d^)-restricted A20 idiotype peptide, CD8^+^ T cells from A20 tumors treated with the eSLC- hCXCL16^K42A^ strain demonstrated increased production of IFNγ and TNFα (**Fig. 2H, S2E**), suggesting increased effector function of tumor antigen-specific T cells following hCXCL16^K42A^ treatment. These data suggest that the eSLC-hCXCL16^K42A^ strain promotes A20 tumor regression via an expansion of activated T cells, specifically tumor antigen-specific T cells.

**Fig. 2.**
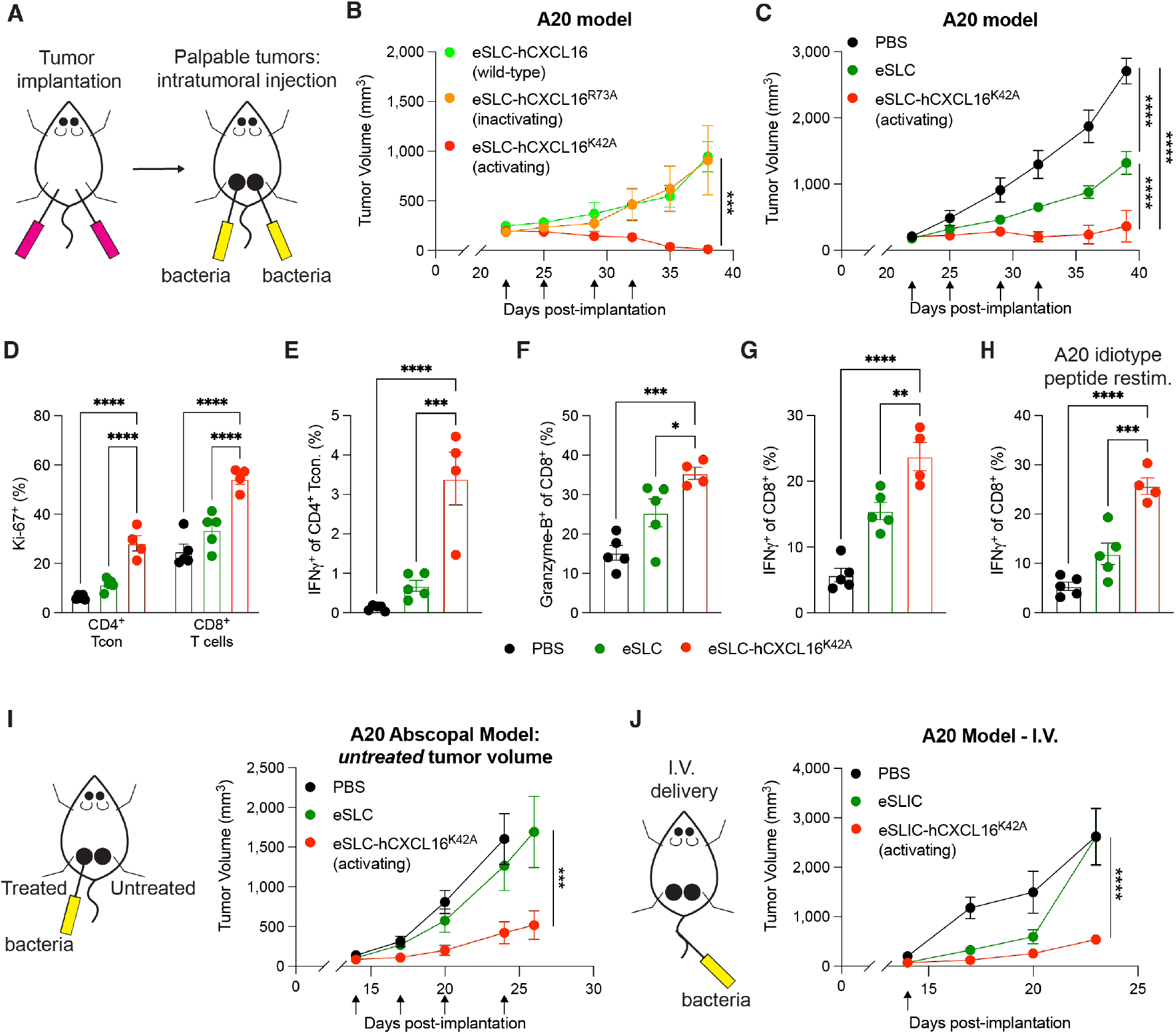
Activating hCXCL16^K42A^ strain promotes tumor regression in subcutaneous murine B cell lymphoma model. **(A)** Schematic overview of implantation and intratumoral treatment of subcutaneously grown A20 B cell lymphoma tumors with EcN strains expressing CXCL16 variants. **(B–C)** Tumor growth curves from BALB/c mice (*n* = 5 per group) subcutaneously implanted with 5 x 10^6^ A20 B cell lymphoma cells on both hind flanks. When tumor volumes were 100–150 mm^3^, mice received intratumoral injections (indicated by black arrows) every 3–4 days with (B) eSLC-hCXCL16, eSLC-hCXCL16^R73A^, or eSLC-hCXCL16^K42A^, or (C) PBS, eSLC, or eSLC-hCXCL16^K42A^. Data are representative of 2 independent experiments (\*\*\**P*<0.001, \*\*\*\**P*<0.0001, 2-way ANOVA with Holm-Sidak post-hoc test at final measurement time point). **(D–H)** Flow cytometric analysis of tumor infiltrating lymphocytes isolated from subcutaneously growing A20 tumors (*n* ≥ 4 mice per group) on day 8 following the second of two intratumoral injections (performed as in (C)) with PBS, eSLC, or eSLC-hCXCL16^K42A^. (D) Frequencies of isolated Ki-67^+^ CD4^+^Foxp3^-^ and CD8^+^ T cells. (E) Frequencies of IFNγ^+^CD4^+^Foxp3^-^ T cells following *ex vivo* restimulation with PMA and ionomycin in the presence of brefeldin A. (F) Frequencies of isolated intratumoral Granzyme-B^+^CD8^+^ T cells. (G) Frequencies of IFNg^+^CD8^+^ T cells following *ex vivo* restimulation with PMA and ionomycin in the presence of brefeldin A. (H) Tumor-infiltrating lymphocytes were stimulated after *ex vivo* isolation with MHC Class I-restricted A20 idiotype peptide (DYWGQGTEL) in the presence of brefeldin A. Frequencies of intratumoral IFNg^+^CD8^+^ T cells. For (D–H), data are representative of 3 independent experiments (\**P*<0.05, \*\**P*<0.01, \*\*\**P*<0.001, \*\*\*\**P*<0.0001, (D) 2-way ANOVA with Holm-Sidak post-hoc test, (E–H) 1-way ANOVA with Holm-Sidak post-hoc test). (I) BALB/c mice (*n* ≥ 5 per group) were implanted subcutaneously with 5 x 10^6^ A20 cells on both hind flanks. When tumor volumes reached 100–150 mm^3^, mice received intratumoral injections every 3–4 days (black arrows) of PBS, eSLC, or eSLC-hCXCL16^K42A^ into a single tumor. Treatment schedule for abscopal assay (left) and tumor growth curves for untreated tumors (right). Data are representative of 2 independent experiments (\*\*\**P*<0.001, 2-way ANOVA with Holm-Sidak post-hoc test at final measurement time point). (J) BALB/c mice (*n* ≥ 5 per group) were injected with 5 x 10^6^ A20 cells into both hind flanks. When tumor volume reached 100–150 mm^3^, mice received a single intravenous injection (black arrow) of PBS, eSLIC, or eSLIC-hCXCL16^K42A^. Schematic for intravenous treatment experiment (left) and tumor growth curves (right). Data are representative of 2 independent experiments (\*\*\*\**P*<0.0001, 2-way ANOVA with Holm-Sidak post-hoc test at final measurement time point). All data displayed as mean ± s.e.m.

To increase the translational relevance, we next aimed to broaden the applicability of this approach beyond accessible subcutaneous tumors. We first assessed the ability of treatment of a single tumor to generate a systemic response to slow growth of untreated distant tumors, termed an ‘abscopal effect’. Using the A20 B cell lymphoma model, we observed that treatment with the hCXCL16^K42A^ strain in one tumor led to slowed tumor growth in distant untreated tumors compared to eSLC alone or PBS (**Fig. 2I, Fig. S2F**). To examine the efficacy of our therapeutics in tumors that cannot be easily accessed for intratumoral injection, we explored intravenous injection as an alternative delivery approach. To this end, we performed a single intravenous injection of bacteria, an approach that specifically colonizes tumors^14,15^, using an EcN strain in which the SLC has been integrated into the genome (eSLIC)^15^. Treatment with eSLIC expressing hCXCL16^K42A^ (eSLIC- hCXCL16^K42A^) significantly reduced the growth of A20 tumors following a single intravenous injection, as compared to treatment with PBS or eSLIC (**Fig. 2J, Fig. S2G**). Taken together, these data demonstrate that hCXCL16^K42A^-expressing EcN strains promote tumor regression, including in distant tumors left untreated and those treated via intravenous injection, and promotes an expansion of activated CD8^+^ T cells.

### Therapeutic efficacy of CXCL16 in murine colorectal and breast cancer models

To assess the broader applicability of this approach, we next examined the therapeutic efficacy of the hCXCL16^K42A^ strain in more aggressive murine cancer models. Treatment of established MC38 colorectal tumors with eSLC-hCXCL16^K42A^ slowed tumor growth compared to PBS and eSLC alone (**Fig. 3A, S3A**). In this colorectal cancer model, we observed an expansion of proliferating conventional CD4^+^ and CD8^+^ T cells following treatment with the hCXCL16^K42A^ strain at day 5 post-treatment (**Fig. 3B**). Furthermore, treatment with the eSLC-hCXCL16^K42A^ led to an expansion of Granzyme-B^+^ CD8^+^ T cells at days 5 and 8 post-treatment (**Fig. 3C-3D**). CXCL16 has been reported to be contribute to anti-tumor immunity in mouse models of triple negative breast cancer (TNBC)^21^. Consistent with this observation, intratumoral injection of eSLC-hCXCL16^K42A^ significantly slowed tumor growth in the TNBC EO771 model compared to PBS or eSLC (**Fig. 3E, S3B**). To examine more translational approaches, we again explored the therapeutic efficacy of intravenous delivery. A single intravenous dose of eSLIC-hCXCL16^K42A^ significantly reduced the growth of MC38 tumors compared to injection of PBS or eSLIC (**Fig. 3F, S3C**). These data show that hCXCL16^K42A^-expressing strains are therapeutically effective, including with a single intravenous treatment, and lead to increased activation of tumor-infiltrating T cells in multiple different murine cancer models, with a more modest benefit in MC38 colorectal and EO771 breast cancer models than in the A20 B cell lymphoma model.

**Fig. 3.**
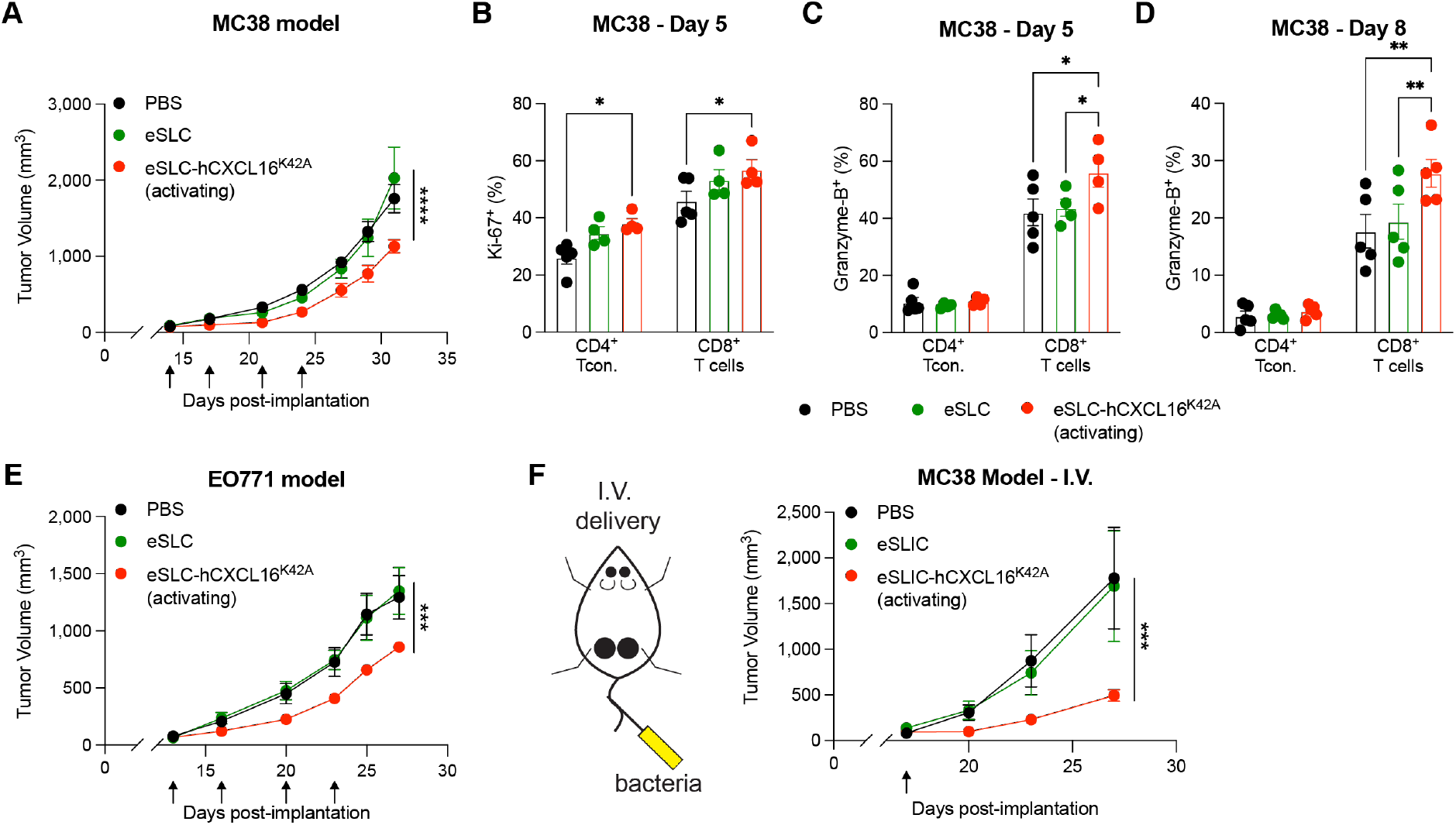
Activating hCXCL16^K42A^ strain offers therapeutic benefit in colorectal and breast cancers. **(A)** Tumor growth curves from C57BL/6 mice (*n* = 5 per group) subcutaneously implanted with 5 x 10^5^ MC38 colorectal tumor cells on both hind flanks. When tumor volumes were 50–150 mm^3^, mice received intratumoral injections (indicated by black arrows) every 3–4 days with PBS, eSLC, or eSLC-hCXCL16^K42A^. Data are representative of 4 independent experiments (\*\*\*\**P*<0.0001, 2-way ANOVA with Holm-Sidak post-hoc test at final measurement time point). **(B–D)** Flow cytometric analysis of tumor infiltrating lymphocytes isolated from subcutaneously growing MC38 tumors (*n* ≥ 4 mice per group) on (B–C) day 5 or (D) day 8 following intratumoral injections (performed as in (A)) with PBS, eSLC, or eSLC-hCXCL16^K42A^. (B) Frequencies of isolated Ki-67^+^ CD4^+^Foxp3” and CD8^+^ T cells. (C–D) Frequencies of isolated Granzyme-B^+^ CD4^+^Foxp3” and Granzyme-B^+^ CD8^+^ T cells on (C) day 5 post-injection and (D) day 8 post-injection. For (B–D), data are representative of 3 independent experiments (\**P*<0.05, \*\**P*<0.01, 2-way ANOVA with Holm-Sidak post-hoc test). **(E)** Tumor growth curves from C57BL/6 mice (*n* = 5 per group) subcutaneously implanted with 1 x 10^6^ EO771 triple negative breast cancer tumor cells on both hind flanks. When tumor volumes were 50–150 mm^3^, mice received intratumoral injections (indicated by black arrows) every 3–4 days with PBS, eSLC, or eSLC-hCXCL16^K42A^. Data are representative of 2 independent experiments (\*\*\**P*<0.001, 2-way ANOVA with Holm-Sidak post-hoc test at final measurement time point). **(F)** C57BL/6 mice (*n* ≥ 5 per group) subcutaneously implanted with 5 x 10^5^ MC38 colorectal tumor cells on both hind flanks. When tumor volume reached 50–150 mm^3^, mice received a single intravenous injection (black arrow) of PBS, eSLIC, or eSLIC-hCXCL16^K42A^. Schematic for intravenous treatment experiment (left) and tumor growth curves (right). Data are representative of 2 independent experiments (\*\*\**P*<0.001, 2-way ANOVA with Holm-Sidak post-hoc test at final measurement time point). All data displayed as mean ± s.e.m.

### Recruitment of dendritic cells synergizes with activated T cell recruitment

As the therapeutic benefit of hCXCL16^K42A^ was more modest in models of colorectal and breast cancer than B cell lymphoma, we explored additional approaches to augment the immune response. T cells are primed by dendritic cells; in particular, type 1 conventional dendritic cells (cDC1) cross-present antigens to activate CD8^+^ T cells. We therefore considered how to recruit dendritic cells to increase T cell priming and thus the availability of activated T cells that could be recruited by treatment with eSLC-hCXCL16^K42A^. Such an approach leverages the unique components of our system: namely, (1) the ability to generate tumor-localized chemokine gradients, and (2) potentiating DC maturation, particularly to type 1 conventional dendritic cells, as enforced by the recognition of bacterial components detected within SLC-generated bacterial lysate. The chemokine CCL20 recruits pre-DCs^22,23^, and when expressed by tumor cells, demonstrates therapeutic potential and dendritic cell recruitment *in vivo*^24,25^. We therefore hypothesized that probiotic-derived CCL20 and CXCL16 would synergize to promote anti-tumor immunity (**Fig. 4A**). Indeed, the combination of hCXCL16^K42A^ and CCL20 strains (“eSLC-combo”) demonstrated a synergistic therapeutic effect in MC38 tumors, slowing tumor growth significantly compared to PBS and eSLC treatment, as well as compared to treatment with either eSLC-hCXCL16^K42A^ or eSLC-CCL20 strains individually (**Fig. 4B, Fig. S4**). Moreover, treatment with the combination of eSLC-hCXCL16^K42A^ and eSLC-CCL20 strains promoted the expansion, both by frequency and number, of type 1 conventional dendritic cells (**Fig. 4C-4D**). Finally, this combination also led to the expansion of tumor-infiltrating, Granzyme-B expressing CD8^+^ T cells (**Fig. 4E**), consistent with increased activation of CD8^+^ T cells by cDC1s.

**Fig. 4.**
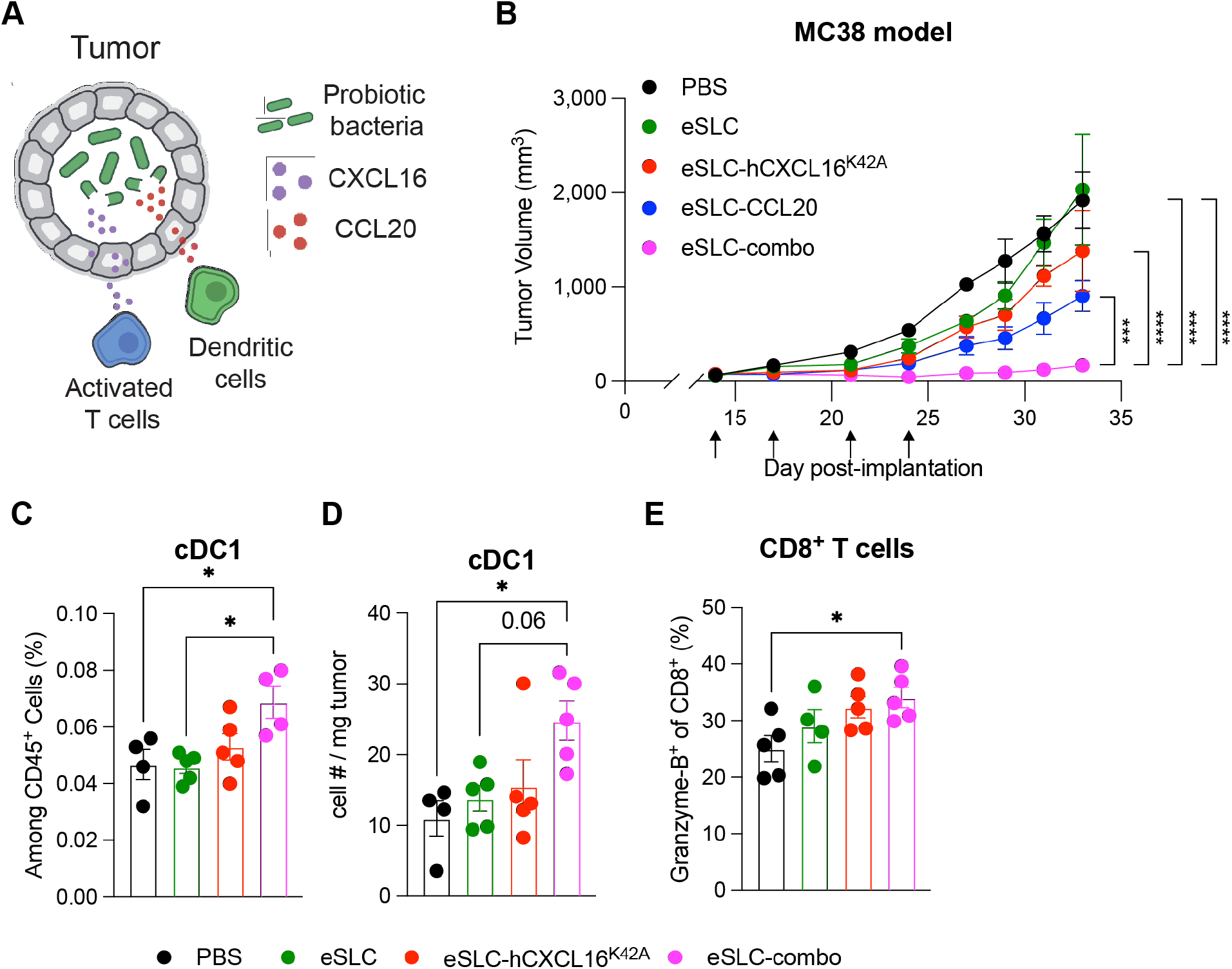
Combination of CXCL16 and CCL20 synergizes to promote anti-tumor immunity. **(A)** Schematic overview of approach to recruit dendritic cells to tumors via CCL20 production in probiotic bacteria to complement activated T cell recruitment via CXCL16 (as in Fig. 1A). **(B)** Tumor growth curves from C57BL/6 mice (*n* = 5 per group) subcutaneously implanted with 5 x 10^5^ MC38 colorectal tumor cells on both hind flanks. When tumor volumes were 50–150 mm^3^, mice received intratumoral injections (indicated by black arrows) every 3–4 days with PBS, eSLC, eSLC-hCXCL16^K42A^, eSLC-CCL20, or eSLC-combo (1:1 mixture of eSLC-hCXCL16^K42A^ and eSLC-CCL20). Data are representative of 4 independent experiments (\*\*\**P*<0.001, \*\*\*\**P*<0.0001, 2-way ANOVA with Holm-Sidak post-hoc test at final measurement time point). **(C–E)** Flow cytometric analysis of tumor infiltrating immune cells isolated from subcutaneously growing MC38 tumors (*n* ≥ 4 mice per group) following intratumoral injections (performed as in (B)) with PBS, eSLC, eSLC-hCXCL16^K42A^, or eSLC-combo (as in (A)). (C) Frequencies and (D) numbers of type 1 conventional dendritic cells on day 5 post-treatment. (E) Frequencies of intratumoral Granzyme-B^+^ CD8^+^ T cells. Data are representative of 3 independent experiments (\**P*<0.05, 1-way ANOVA with Holm-Sidak post-hoc test). All data displayed as mean ± s.e.m.

## DISCUSSION

Here we demonstrate the utility of recruiting cells responsible for the adaptive immune response into tumors. We find that expression of hCXCL16^K42A^ recruits activated mouse T cells, offers therapeutic benefit *in vivo*, and leads to the expansion of effector T cells in the tumor. Furthermore, we observe synergy between the recruitment of dendritic cells and activated T cells in slowing tumor growth. We believe this system offers a new set of therapeutic strategies to elicit immune cell recruitment and may complement existing immunotherapies that target the activation of immunity systemically but are unable to act locally to increase the intratumoral infiltration of cells.

Bacteria-derived CXCL16 alone has substantial therapeutic efficacy, and the use of an activating mutation of human CXCL16 that is bioactive on mouse and human T cells in our report may reduce translational studies required to show therapeutic pre-clinical benefits. CXCL16 alone does not promote tumor regression in all of the murine tumor models assessed, with one possible explanation being a paucity of activated T cells. In this regard, the combination of CXCL16 with CCL20 shows promise for addressing the possible lack of activated T cells in certain tumor types. Our study focuses on the chemokines CXCL16 and CCL20, but other chemokines, or combinations of chemokines, may offer additional therapeutic benefit. Our approach also has advantages over engineering tumor cells to express chemokines using oncolytic viruses and other gene delivery approaches due to the simplicity of bacterial growth and delivery to the tumor microenvironment, either by intratumoral or intravenous injection, with repeated release of therapeutic payloads occurring through multiple growth and lysis cycles.

Although checkpoint blockade offers durable survival in previously untreatable disease^26^, it only achieves disease regression in a small subset of patients, with immunologically ‘hot’ tumors offering greater responsiveness compared to ‘cold’ tumors^2–4^. Indeed, the recruitment of peripheral tumor-specific T cells may be one mechanism to enhance the therapeutic response of PD-1/PD-L1 blockade^27,28^. Furthermore, the effectiveness of tumor vaccines and cell therapies, such as *ex vivo* expanded tumor infiltrating lymphocytes and chimeric antigen receptor T cells, is hindered by the inability of primed T cells to infiltrate tumors^29,30^. Our approach may therefore offer a means to overcome challenges in other immunotherapies, broadening their efficacy and applicability. In summary, we show effectiveness of engineered bacteria to recruit activated T cells and dendritic cells to tumors, which results in slowed tumor growth and increased effector T cell and dendritic cell infiltration. This approach offers a new range of therapeutic strategies in the quest for novel anti-cancer therapies.

## METHODS

### Bacterial strains

The mature bioactive regions of murine (Asn27-Pro114, UniProt Accession Number Q8BSU2) and human (Asn30-Pro118, UniProt Accession Number Q9H2A7) CXCL16, and murine CCL20 (Ala27-Met96, UniProt Accession Number O89093) were cloned into plasmid p246^15^ via Gibson Assembly, (co)transformed with or without the SLC plasmid (p15a) into electrically competent *E. coli* Nissle 1917 (EcN), and maintained as previously described^15^. Briefly, strains with only a therapeutic plasmid (p246) were grown in LB broth with kanamycin (50 μg/mL). Strains with the therapeutic and SLC plasmids were grown in LB broth with kanamycin (50 μg/mL) and spectinomycin (100 μg/mL) with 0.2% glucose. Mutant human CXCL16^K42A^ and CXCL16^R73A^ plasmids were generated from the wildtype hCXCL16 p246 plasmid by the New England BioLabs Q5 Site-Directed Mutagenesis Kit, as per manufacturer’s instructions.

### Chemotaxis assay

Activated T cells were generated as described^16^. Briefly, T cells were isolated from wildtype adult C57BL/6 mouse spleen and lymph nodes using the Dynabeads FlowComp Mouse Pan T (CD90.2) Kit (Thermo Fisher Scientific) as per the manufacturer’s protocol. Isolated T cells were cultured with anti-CD3/CD28 beads (Dynabeads Cat. # 11452D; Thermo Fisher Scientific) in a 1:1 ratio in 10% complete RPMI (RPMI 1640 medium supplemented with 10% FBS, Pen/Strep, non-essential amino acids, Glutamax, HEPES, Sodium Pyruvate and 2-Mercaptoethanol). After 5 days, the beads were removed by magnetic separation and T cells were replated at 10^6^ cells/mL for 4 days in 10% complete RPMI supplemented with 100 IU/mL of rhIL-2. T cells were then washed and resuspended at 5.9×10^6^ cells/mL in serum-free complete RPMI in preparation for the chemotaxis assay. Human T cells from anonymous donors sourced by STEMCELL Technologies were isolated by negative selection for CD3^+^ populations as per manufacturer’s instruction (EasySep, STEMCELL Technologies), and cultured and prepared identically to murine T cells using corresponding reagents for human cells.

Overnight cultures of each bacterial strain (without SLC) were grown in LB with appropriate antibiotics then subcultured at a 1:100 dilution in a shaking incubator for 90 min at 37 °C. Bacteria were washed twice in serum-free complete RPMI, OD600 matched, and lysed via sonication in serum-free complete RPMI. Lysates were centrifuged to remove debris (20,817xg for 10 min at 4°C) and 235 μL of the supernatant was entered into the lower chamber of a Corning HTS transwell plate (well area = 0.143 cm^2^; pore size = 5 μm). Activated T cells (75 μL of the preparation described above) were added to the upper chamber and the plate was incubated for 3 hours in a humidified 37 °C 5% CO2 incubator. The bottom chamber was then harvested, washed, and stained with anti-mouse CD3 violetFluor450 (Tonbo Biosciences; clone 17A2), CD4 APC (Tonbo Biosciences; clone RM4-5) and CD8 PE (Tonbo Biosciences; clone 53-6.7). Human T cells were stained with anti-human CD3 BUV395 (BD Biosciences; clone SK7), CD4 BV785 (BioLegend; clone RPA-T4), and CD8 BV510 (BD Biosciences; clone RPA-T8). Samples were then acquired on a BD Fortessa for 60 seconds. Cell counts were normalized to the number of cells entered into the assay.

### Human CXCL16 ELISA

For *in vitro* characterization, relevant strains were grown overnight as described above then subcultured (1:100 dilution) for 3 hours in a shaking incubator at 37 °C. Cultures were then centrifuged (20,817xg for 10’ at 4 °C) and supernatants were entered into a hCXCL16 ELISA (R&D Human CXCL16 DuoSet, Cat. # DY1164), performed per the manufacturer’s protocol. For quantification of hCXCL16 concentrations in tumor homogenates, three days after the final intratumoral injection of bacteria, tumors were harvested, weighed, and homogenized in tissue lysis buffer (100 mM Tris pH=7.4, 150 mM NaCl, 1 mM EGTA, 1 mM EDTA, 1% Triton-X100, 0.5% sodium deoxycholate in water) with protease inhibitor and EDTA. Homogenates were then centrifuged (5000xg for 10’ at 4 °C) to remove debris and supernatants were used for ELISA, performed per the manufacturer’s protocol.

### Animal models

All animal experiments were approved by the Institutional Animal Care and Use Committee of Columbia University (protocols AC-AAAQ8474, AC-AABD8554, and AC-AAAZ4470). The protocol requires animals to be euthanized when the tumor burden reaches 2 cm in diameter or upon recommendation by veterinary staff. A20 cells were maintained in RPMI supplemented with 10% FBS, Pen/Strep and 2-Mercaptoethanol. MC38 and EO771 cells were maintained in DMEM supplemented with 10% FBS, Pen/Strep, non-essential amino acids, Glutamax, HEPES, Sodium Pyruvate and 2-Mercaptoethanol. Cultures were maintained in a humidified 37 °C, 5% CO2 incubator. Prior to injection, A20 cells were resuspended in in RPMI without phenol red at 5 x 10^7^ cells/mL. A20 cells were subcutaneously implanted at 100 μL (5 x 10^6^ cells) per hind flank. MC38 and EO771 cells were washed in PBS, resuspended at 5 x 10^6^ cells/mL and 10 x 10^6^ cells/mL in PBS, respectively, and 100 μL of cell suspension was then injected subcutaneously into both hind flanks. Female 7-8 week old BALB/c (for A20 tumors) or C57BL/6N (EO771 and MC38 tumors) mice were purchased from Taconic Biosciences or Jackson Laboratories, allowed to acclimate for a week and then injected with tumor cells. A20 tumor volume was determined by caliper measurements (length x width^2^ x 0.5) and mice were randomly assigned to treatment groups after tumors reached a volume of 100–150 mm^3^. MC38 and EO771 tumor volume was calculated as length x width x height, and mice were randomly assigned to treatment groups after tumors reached a volume of 50–150 mm^3^. For treatment with bacteria, bacteria were cultured in a 37 °C shaking incubator for up to 12 hours to reach stationary phase of growth in LB broth with appropriate antibiotics and 0.2% glucose. Bacteria were then subcultured (1:100 dilution) until a maximum OD_600_ of 0.15 was reached, again in LB broth with appropriate antibiotics and 0.2% glucose, then washed 3 times and resuspended in ice cold PBS. A total of 40 μL of each suspension containing 5 x 10^6^ (for A20 tumors) or 1 x 10^6^ bacteria (for MC38 and EO771 tumors) were injected intratumorally. Intratumoral injections were performed every 3–4 days for a total of 3–4 treatments (for tumor growth assays), or 1–2 treatments (for immunophenotyping assays), as indicated. For intravenous treatment experiments, bacteria were prepared as above and injected via the tail-vein at a concentration of 5 x 10^7^ per mL in PBS, with a total volume of 100 μL injected per mouse.

### Immune phenotyping by flow cytometry

Tumors were treated as above and harvested at the indicated time points. For isolation of tumor infiltrating lymphocytes, tumors were excised, then minced and digested in wash media (RPMI 1640 supplemented with 5% FCS, HEPES, Glutamax, Pen/Strep) with 1 mg/mL collagenase A and 0.5 μg/mL DNAse I in a shaking incubator for up to 45 minutes at 37 °C to achieve a single cell suspension. Once a single cell suspension was achieved, samples were washed, then either restimulated or stained for flow cytometry analysis. For cytokine staining and *ex vivo* restimulation with PMA and ionomycin, aliquots of tumor homogenates were incubated for 3 hours at 37 °C in 10% complete RMPI (as above) with PMA (50 ng/mL), ionomycin (500 ng/mL) and brefeldin A (1 μg/mL) prior to flow cytometry staining. For cytokine staining and *ex vivo* restimulation with A20 idiotype peptide, aliquots of tumor homogenates were incubated for 5 hours at 37 °C in 10% complete RMPI (as above) with the A20 idiotype peptide (DYWGQGTEL; 1 μg/mL) and brefeldin A (1 μg/mL) prior to staining for flow cytometry. Live/dead staining was performed via Ghost Dye Red 780 labeling (Tonbo Biosciences), as per the manufacturer’s protocol. Cells were then stained for flow cytometry, with intracellular staining performed using the Tonbo Foxp3/Transcription Factor Staining Buffer Kit per manufacturer’s instructions. Antibodies used were anti-CD45 (clone 30-F11, BioLegend), NK1.1 (clone PD136, BD Biosciences), CD3e (clone 145-2C11, Tonbo Biosciences), TCRß (clone H57-597, BD Biosciences), CD4 (clone RM4-5, BD Biosciences), CD8 (clone 53-6.7, Tonbo Biosciences), Foxp3 (clone FJK-16s, Thermo Fisher Scientific), Granzyme-B (clone QA16A02, BioLegend), Ki-67 (clone SolA15, Thermo Fisher Scientific), IFNγ (clone XMG1.2, Tonbo Biosciences), B220 (clone RA3-6B2, BD Biosciences), CD11c (clone N418, Tonbo Biosciences), Ly6G (clone 1A8, Tonbo Biosciences), CD11b (clone M1/70, Tonbo Biosciences), MHCII (clone M5/114.15.2, Tonbo Biosciences), CD103 (clone 2E7, BioLegend).

## Acknowledgements

We thank members of the Arpaia and Danino labs for helpful discussions and members of the S.H. Sternberg lab for technical advice. This work was supported by NIH/NCI R01CA249160 (N.A. and T.D.), NIH/NCI R01CA259634 (N.A.), Searle Scholars Program SSP-2017-2179 (N.A.), the Roy and Diana Vagelos Precision Medicine Pilot Grant (N.A. and T.D.), and NIH/NIGMS T32GM007367 (T.M.S.). Research reported in this publication was performed in the Columbia University Department of Microbiology & Immunology Flow Cytometry Core facility. The content is solely the responsibility of the authors and does not necessarily represent the official views of the National Institutes of Health.

## Contributions

T.M.S. and N.A. conceived and designed the project with input from T.D.; T.M.S. performed experiments and analyzed data jointly with N.A.; R.L.V. assisted with transwell assays, S.S.R., A.A. and F.L. assisted with tumor immunophenotyping, L.H.H., K.P., and C.C. assisted with a portion of the animal studies. T.M.S. and N.A. wrote the manuscript.

---

**Fig. S1.**
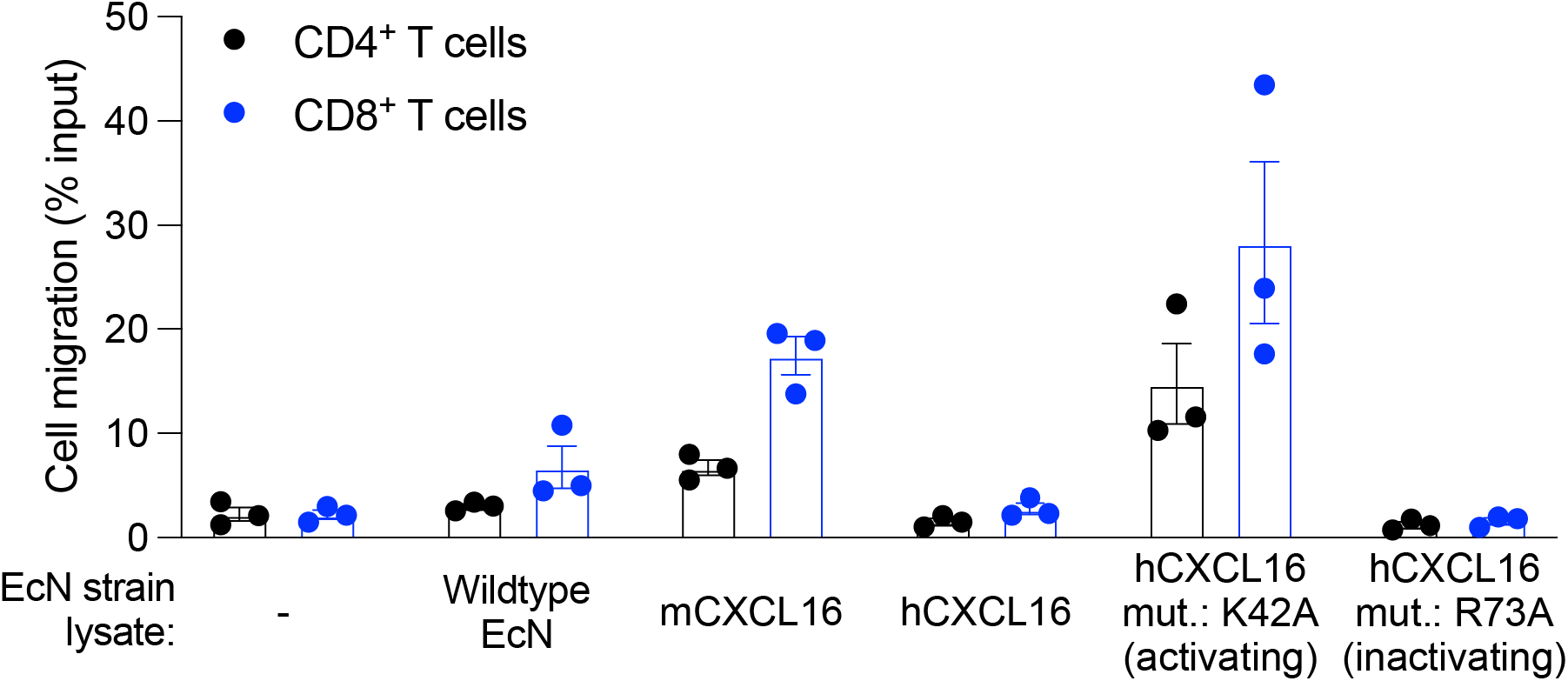
Activating mutation of human CXCL16 has bioactivity on activated human T cells. Flow cytometric quantification of the migration of activated human T cells in response to each of the indicated EcN strain lysates. Media without any bacteria was used as a control (“-”) Representative of 2 independent human controls. All data displayed as mean ± s.e.m.

**Fig. S2.**
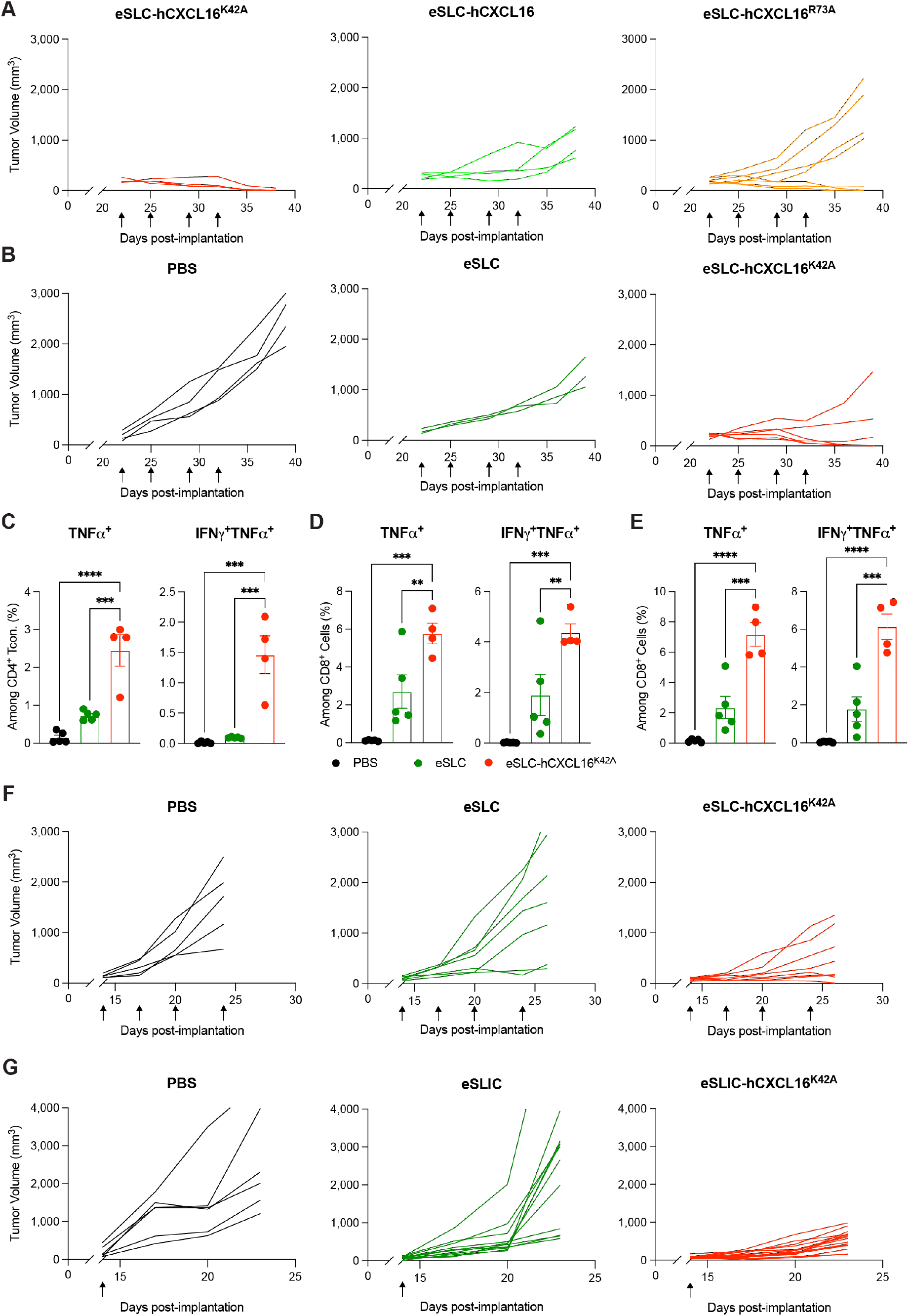
Activating mutation of human CXCL16 in syngeneic A20 tumors. **(A-B)** Individual tumor growth curves from BALB/c mice (*n* = 5 per group) subcutaneously implanted with 5 x 10^6^ A20 B cell lymphoma cells on both hind flanks. When tumor volumes were 100–150 mm^3^, mice received intratumoral injections (indicated by black arrows) every 3–4 days with (A) eSLC- hCXCL16, eSLC-hCXCL16^R73A^, or eSLC-hCXCL16^K42A^, or (B) PBS, eSLC, or eSLC- hCXCL16^K42A^. Data are representative of 2 independent experiments. **(C-E)** Flow cytometric analysis of tumor infiltrating lymphocytes isolated from subcutaneously growing A20 tumors (*n* ≥ 4 mice per group) on day 8 following the second of two intratumoral injections (performed as in (B)) with PBS, eSLC, or eSLC-hCXCL16^K42A^. (C) Frequencies of TNFα^+^ CD4^+^Foxp3^-^ or TNFα^+^IFNγ^+^CD4^+^Foxp3^-^ T cells and (D) Frequencies of TNFα^+^ CD8^+^ or TNFα^+^IFNγ^+^CD8^+^ T cells following *ex vivo* restimulation with PMA and ionomycin in the presence of brefeldin A. (E) Tumor-infiltrating lymphocytes were stimulated after *ex vivo* isolation with MHC Class I-restricted A20 idiotype peptide (DYWGQGTEL) in the presence of brefeldin A. Frequencies of intratumoral TNFα^+^ CD8^+^ or TNFα^+^IFNγ^+^CD8^+^ T cells. Data are representative of 3 independent experiments (\*\**P*<0.01, \*\*\**P*<0.001, \*\*\*\**P*<0.0001, 1-way ANOVA with Holm-Sidak post-hoc test). Data displayed as mean ± s.e.m. **(F)** BALB/c mice (*n* ≥ 5 per group) were implanted subcutaneously with 5 x 10^6^ A20 cells on both hind flanks. When tumor volumes reached 100– 150 mm^3^, mice received intratumoral injections every 3–4 days (black arrows) of PBS, eSLC, or eSLC-hCXCL16^K42A^ into a single tumor. Individual tumor growth curves for untreated tumors. Data are representative of 2 independent experiments. **(G)** BALB/c mice (*n* ≥ 5 per group) were injected with 5 x 10^6^ A20 cells into both hind flanks. When tumor volume reached 100–150 mm^3^, mice received a single intravenous injection (black arrow) of PBS, eSLIC, or eSLIC- hCXCL16^K42A^. Individual tumor growth curves displayed. Data are representative of 2 independent experiments.

**Fig. S3.**
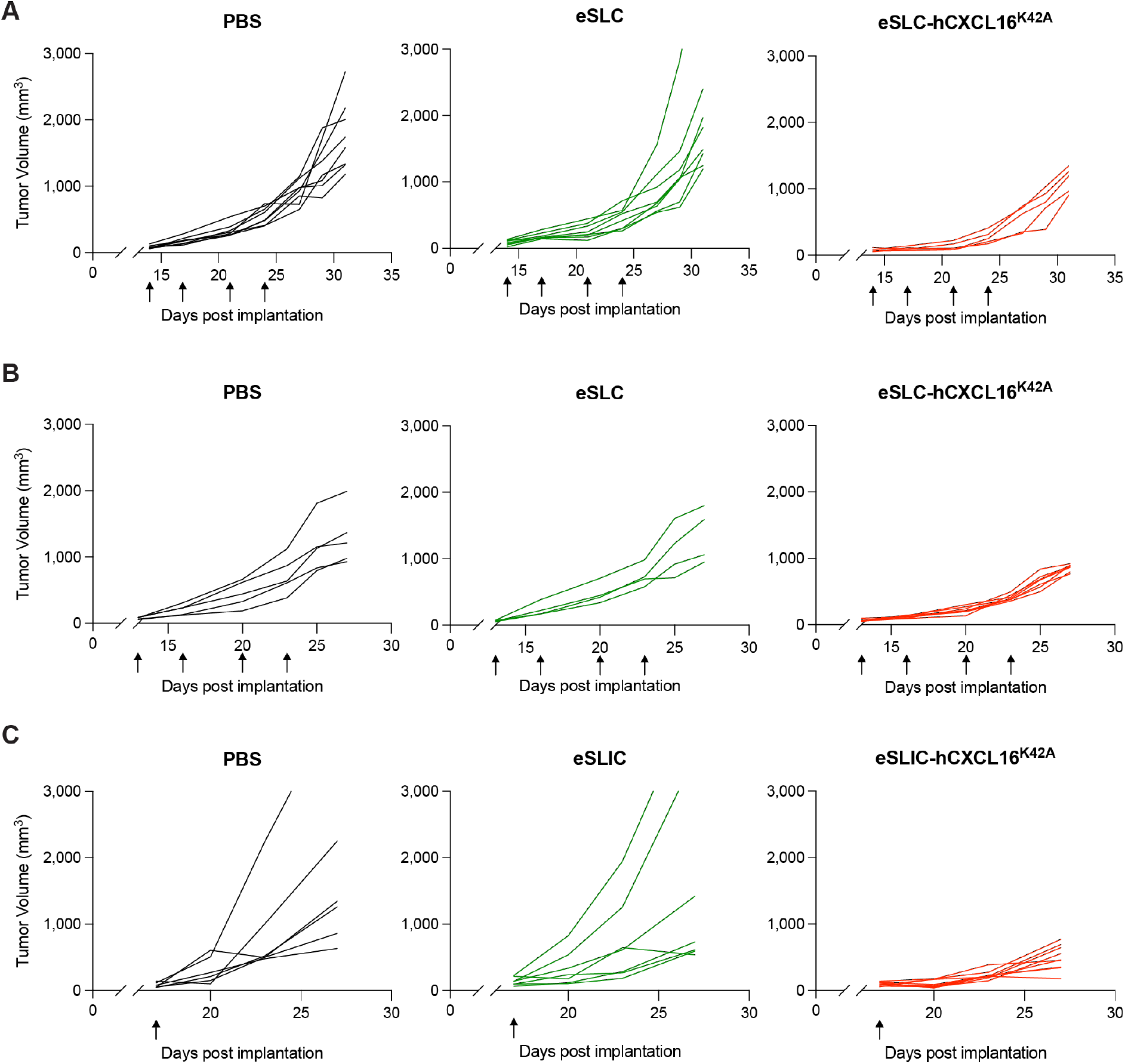
Individual tumor trajectories of syngeneic tumors treated with CXCL16 or controls. **(A-B)** Individual tumor growth curves from C57BL/6 mice (*n* = 5 per group) subcutaneously implanted with (A) 5 x 10^5^ MC38 colorectal tumor cells or (B) 1 x 10^6^ EO771 triple negative breast cancer cells on both hind flanks. When tumor volumes were 50–150 mm^3^, mice received intratumoral injections (indicated by black arrows) every 3–4 days with PBS, eSLC, or eSLC- hCXCL16^K42A^. (A): Data are representative of 4 independent experiments. (B) Data are representative of 2 independent experiments. **(C)** C57BL/6 mice (*n* ≥ 5 per group) subcutaneously implanted with 5 x 10^5^ MC38 colorectal tumor cells on both hind flanks. When tumor volume reached 50–150 mm^3^, mice received a single intravenous injection (black arrow) of PBS, eSLIC, or eSLIC-hCXCL16^K42A^. Individual tumor growth curves displayed. Data are representative of 2 independent experiments.

**Fig. S4.**
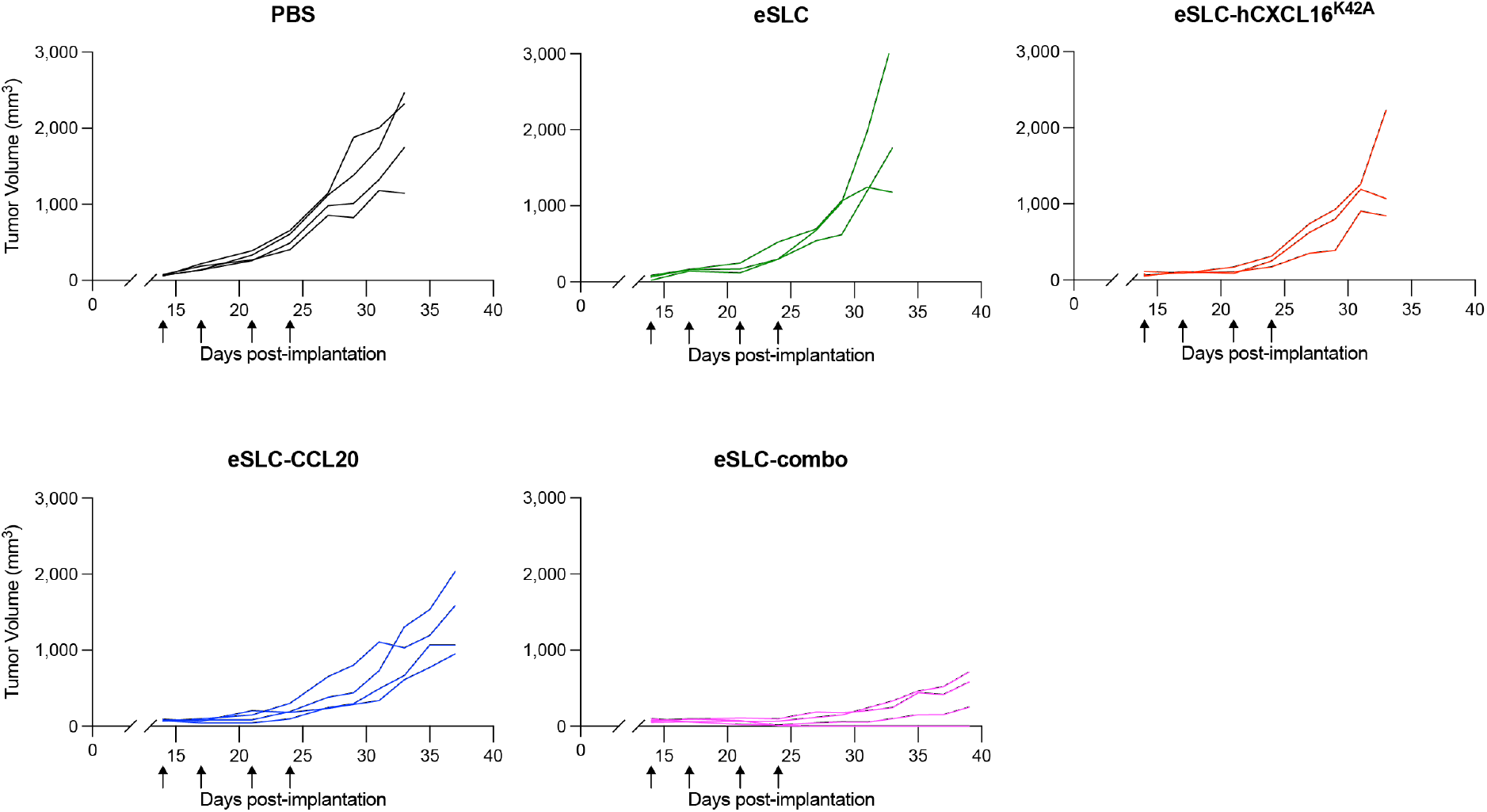
Individual tumor trajectories of MC38 tumors in combination therapy approach. Individual tumor growth curves from C57BL/6 mice (*n* = 5 per group) subcutaneously implanted with 5 x 10^5^ MC38 colorectal tumor cells on both hind flanks. When tumor volumes were 50–150 mm^3^, mice received intratumoral injections (indicated by black arrows) every 3–4 days with PBS, eSLC, eSLC-hCXCL16^K42A^, eSLC-CCL20, or eSLC-combo (1:1 mixture of eSLC-hCXCL16^K42A^ and eSLC-CCL20). Data are representative of 4 independent experiments.

